# PolySTest: Robust statistical testing of proteomics data with missing values improves detection of biologically relevant features

**DOI:** 10.1101/765818

**Authors:** Veit Schwämmle, Christina E Hagensen, Adelina Rogowska-Wrzesinska, Ole N. Jensen

**Author notes:** **Abbreviations:** *FDR*, False discovery rate, *DRF*, Differentially regulated feature.

## Abstract

Statistical testing remains one of the main challenges for high-confidence detection of differentially regulated proteins or peptides in large-scale quantitative proteomics experiments by mass spectrometry. Statistical tests need to be sufficiently robust to deal with experiment intrinsic data structures and variations and often also reduced feature coverage across different biological samples due to ubiquitous missing values. A robust statistical test provides accurate confidence scores of large-scale proteomics results, regardless of instrument platform, experimental protocol and software tools. However, the multitude of different combinations of experimental strategies, mass spectrometry techniques and informatics methods complicate the decision of choosing appropriate statistical approaches. We address this challenge by introducing PolySTest, a user-friendly web service for statistical testing, data browsing and data visualization. We introduce a new method, Miss Test, that simultaneously tests for missingness and feature abundance, thereby complementing common statistical tests by rescuing otherwise discarded data features. We demonstrate that PolySTest with integrated Miss Test achieves higher confidence and higher sensitivity for artificial and experimental proteomics data sets with known ground truth. Application of PolySTest to mass spectrometry based large-scale proteomics data obtained from differentiating muscle cells resulted in the rescue of 10%-20% additional proteins in the identified molecular networks relevant to muscle differentiation. We conclude that PolySTest is a valuable addition to existing tools and instrument enhancements that improve coverage and depth of large-scale proteomics experiments. A fully functional demo version of PolySTest and Miss Test is available via http://computproteomics.bmb.sdu.dk/Apps/PolySTest.

## Introduction

A typical mass spectrometry based proteomics study identifies and quantifies thousands of proteins in large scale experiments, including different perturbations, cell types or organisms, dose-responses or time courses. Experimental and biological variance is captured by analyzing technical and biological replicates in order to enhance reproducibility and repeatability. However, the extensive resource requirements of experimental, large-scale proteomics experiments usually result in low replicate numbers, i.e. *n* = 2 − 5. This limitation requires careful application of appropriate statistical methods. Given the measured values of thousands of peptides or proteins, biological interpretation of the large data sets calls for data processing and filtering for detection of the biologically relevant features. This is usually achieved by the detection of differentially regulated features (DRFs), such as, differentially expressed proteins, peptides or post-translationally modified peptides. Statistical testing, also known as significance analysis, helps identifying biological changes in the experimental setup. Statistical tests aim to recognize DRFs by providing probabilities for a significant change of protein abundance. Here, variance between biological samples, technical variation and “computational variance” coming from applying a particular data processing workflow [1, 2] determines the significance. Optimal application of suitable statistical tests relies on estimating these variances to yield the correct false discovery rate (FDR, defined by the number of false positives divided by the number of DRFs) while maximizing the number of correctly identified proteins (sensitivity).

Particularly “missing values” can compromise statistical testing as they severely decrease the statistical power of the data analysis. Missing values are values of a feature that are absent in a given replicate by not having been detected and reported by the measurement equipment. A missing value originates because a signal is below the detection limit, due to sample loss, and/or stochastic precursor selection in mass spectrometry. The different origin of missing values in a data set impedes accurate prediction of the correct values. Thus, imputation methods that replace missing values with estimated abundances can be considered inappropriate as they add knowingly false measurements. Even when assuming missing values to be due to absent proteins, applying imputation by 0-values would lead to an estimated variance of zero and impede transforming the values to their logarithm. Therefore, data imputation has been reported to lead to erroneous results [3, 4]. Currently, most computational approaches apply a prior filter to exclude proteins of low coverage that are missing too many values and do not provide sufficient replication to estimate their variance by having only 1-2 values per experimental condition.

However, proteins that are represented in datasets with both “present values” and “missing values” still contain highly valuable information. Missing values coming from proteins of low or no abundance can contribute to a statistical test with additional evidence. For instance, consider a case where a protein is completely repressed and thus all replicate data are missing in one condition, but are highly abundant in the other condition. Including these otherwise discarded cases using a statistically sound method would rescue more proteins for the statistical testing and so is likely to increase the number of differentially regulated features (DRFs). Current tests with capability to include missing values however are usually binary, i.e. they oversimplify the analysis by dividing all data values into absent and present, or by assuming all missing values to be below the detection limit [5, 6].

The fraction of missing values often increases drastically on the peptide level. Peptide-centric approaches where the statistical test includes peptide quantifications instead of summarized protein amounts are very promising. They extrapolate protein variance from peptide variance and some tools are already available, including MSStats [7] and MSqRob [4]. However, in most cases rather low numbers of quantified peptides per protein complicate the performance of such approaches. In addition, different mass spectrometry data acquisition methods (e.g. label-free versus stable isotope labeling protocols) require specialized methods for protein summarization. Therefore, we here consider summarization as a separate task.

Several methods can process summarized data with few missing values and produce satisfactory results. The LIMMA method [8], originally developed for micro-array data, is widely used in proteomics research. LIMMA was shown to perform well for proteomics experiments with as little as three replicates per condition [9]. However, depending on the particular data set and its structure, other statistical tests such as rank products [10] were found to improve performance in a complementary manner. Distinct data properties originating from different numbers of differentially regulated features and different variations within them lead to either rank products or LIMMA becoming the best-performing method in ground truth data [9]. Another broadly used approach that holds potential to deal with differently structured data is based on comparing the observed values to randomizations of the data. Such permutation tests have been implemented e.g. in the Perseus suite [11].

Given the current state of available methods, there is a need to carry out significance analysis that a) considers missing values, b) uses robust and complementary tests to include different intrinsic data properties and c) provides simple usage through a user-friendly program interface.

We here introduce a novel statistical test (Miss Test) for datasets with missing values. The Miss Test is included in our new PolySTest web service that provides a set of complementary statistical tests for quantitative data and versatile data browsing and visualization. Validation of the methods by extensive tests with artificial and real data sets shows the power of our approach. We demonstrate the performance of PolySTest using a proteomics data set to achieve improved coverage of biologically significant features in muscle cell differentiation.

## Methods

### Missing value statistics (Miss test)

The probability *p*_NA_ to find a missing value is given by the fraction of missing values in the data set.

A binomial distribution describes the case of having multiple missing measurements. The probability to find *i* missing values in *r* replicates is given by

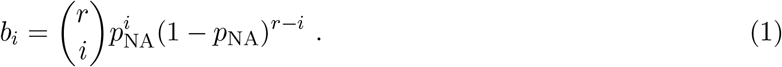

We derive the probability to observe a difference of the number of missing values in two groups of values.

Then the probability for finding a difference of *k* missing values between sample groups yields

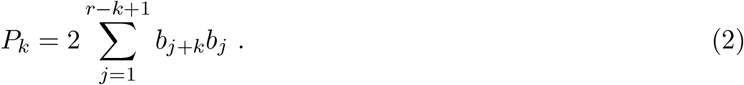

For an example for calculating the probabilities, see the Result section.

The probabilities *P*_*k*_ only account for presence/absence of protein quantifications but not for their abundance-dependence, i.e. whether they are more likely to occur when measuring low abundant proteins. Therefore, the algorithm applies the following additional steps. The distribution of all quantitative values is divided into 100 quantiles. Then, for each quantile, a) values in the quantiles below the selected one are removed; b) probabilities described above (Eq. 1) are calculated on basis of the new *p*_NA_; c) the difference in number of missing values between conditions for each feature is taken; and d) the probabilities for the differences using Eq. 2 are stored. This method gives, for each feature and each comparison between conditions, a vector of 100 *P*_*k*_ values. The smallest *P*_*k*_ of each vector is multiplied by *r* + 1 and represents the p-value of the Miss test.

### Data sources

#### Artificial data sets

Real data with ground truth was simulated by superposing a fraction of the normally distributed (mean 0, standard deviation 1) features with an offset. Out of *N* features per replicate (*R*) and one of the two conditions, *N*_*R*_ features were displaced by *δ* to each side with random assignment of up- and down-regulations.

Then, for the simulation of abundance-dependent missingness, we randomly removed *m*% of the values by elimination with weights [1 − *r*(*i*)*/N*]^*µ*^ where *r*(*i*) is the rank of the feature *i* in a sample after sorting for their abundance. Table 1 shows a summary of the parameter ranges used for creating the artificial data sets.

**Table 1:** Parameter values of artificial data including their range.

| Parameter | Symbol | Range |
| --- | --- | --- |
| Feature number | $N$ | 500-10,000 |
| Replicates per condition | $R$ | 3-10 |
| Percentage regulated features | $N_R$ | 0-50% |
| Difference regulated features | $\delta$ | 1-5 |
| Percentage missing values | $m$ | 0-50% |
| Abundance-dependence of missing values | $\mu$ | 0-100 |

#### Experimental data sets

##### 3-species mixture

The original experiment [12] comprises two hybrid proteome samples with a “ground truth” mix of human, yeast and Escherichia coli proteins (ratios 1:1, 2:1 and 1:4, respectively), referred to as HYE124. The samples were analyzed in technical triplicates on a TripleTOF 6600 instrument using a 64 variable window of 10 min for SWATH-MS acquisition. The protein quantification summary from data analysis with SWATH 2.0 was chosen. The file `PViewBuiltinProteins_TTOF6600_64w_shift_iRT_extractionWindo` was downloaded from PRIDE repository (PXD002952) and the protein quantification values were log-transformed.

##### 2-species mixture

We retrieved data from a study where Escherichia coli digest was spiked into a HeLa digest in four different concentrations: 3%, 7.5%, 10% and 15% [13]. The data was acquired via label-free MS on a Q Exactive Plus instrument. Protein abundances were given as supplementary data file. These were log-transformed and normalized using the normalizeCyclicLoess function (LIMMA R package [14]).

##### Muscle differentiation

Immortalized human satellite cells (KM155C25) [15] were differentiated into muscle cells and samples. Biological triplicates were taken at 6 time points: proliferating activated myoblasts (Day −1) and 5 days following the initiation of differentiation into mature myocytes (Day 0, 1, 2, 3 and 4). The data was acquired using an Orbitrap Fusion™Lumos™Tribrid™Mass Spectrometer. Protein quantifications were taken from MaxQuant and contaminants and decoy hits were removed. Quantifications from technical replicates were summed, log_2_-transformed and normalized by median. Furthermore a minimum of 2 unique peptides per protein and 6 non-missing values over all 18 samples was required to reduce the effect of wrong identifications.

### Statistical tests

*t-test* Paired or unpaired t-tests were carried out using the default *t.test* in R. Resulting p-values were corrected for multiple testing according to Storey [16] resulting in FDR estimations.

*LIMMA* Paired or unpaired moderated tests were calculated applying the default procedure for using the limma package [14] including linear model estimation and empirical Bayes moderation of standard errors. Correction for multiple testing was again obtained according to Storey.

*rank products* The p-values for the paired tests were calculated according to ref. [17]. Unpaired samples were simulated by obtaining the p-values from 100 randomly picked pairings and taking their means. Correction for multiple testing was by Benjamini-Hochberg because Storey was not found to be sufficiently robust in some test cases.

*permutation test* This method requires a minimum of 7 replicates per condition (unpaired) or 7 ratios between pairings. For lower replicate numbers, the method adds samples with values randomly drawn from the pool containing all values of the considered samples. Then, for each feature, *u*_paired_ or *u*_unpaired_ were obtained for 1,000 randomizations of the feature values. Similar to calculating the t-values in a t-test, the mean of all paired ratios was divided by their standard deviation. For feature *i*, this gives 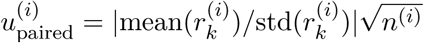 where 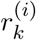 is the abundance of feature *i* in replicate *k* and *n*^(*i*)^ is the number of non-missing values. Unpaired comparison of values of feature *i* between conditions *t* and *c* was calculated by

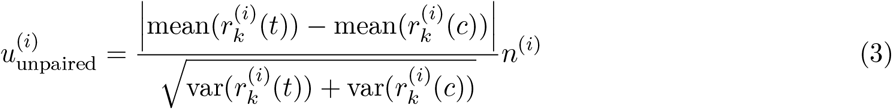

where lower replicate numbers are punished by multiplying by the number of non-missing values.

We then calculated the p-value of each feature by comparing *u*^(*i*)^ of the feature to 1,000 instances of *u*^(*r*)^ that were calculated for the randomizations *r* explained above. The resulting p-value is given by

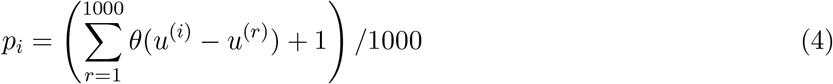

where *θ*(*x*) is the Heaviside function which is 1 for *x >* 0 and 0 otherwise.

Correction for multiple testing by the Benjamini-Hochberg provides false discovery rates.

*Miss test* For details on this statistical test, see above.

*Pathway enrichment analysis* was carried out using the ClusterProfiler R package [18] (version 3.10.1) using default parameters.

### Web interface

All methods are freely available through our web service at http://computproteomics.bmb.sdu.dk/Apps/PolySTest or can be downloaded to be run on a local computer. The software was written as Shiny app which is highly interactive and allows extensive data browsing and visualization including upset plots [19], circlize plots [20] and an interactive heatmap [21]. The full source code is available through https://bitbucket.org/veitveit/polystest.

## Results

We introduce a new statistical method, Miss test, to combine the occurrence of missing values with protein abundance for improved detection of DRFs in data with high amounts of missing values. To assess its performance, robustness to different data structures and complementarity to other statistical tests, we assessed five different statistical tests applied on hundreds of artificial data sets, experimental data sets with ground truth and a data set with well-defined biological content.

The tests are implemented and combined in the PolySTest web service to analyze and visualize the statistically tested data prior to down-stream biological interpretation (Fig 1A).

**Figure 1:**
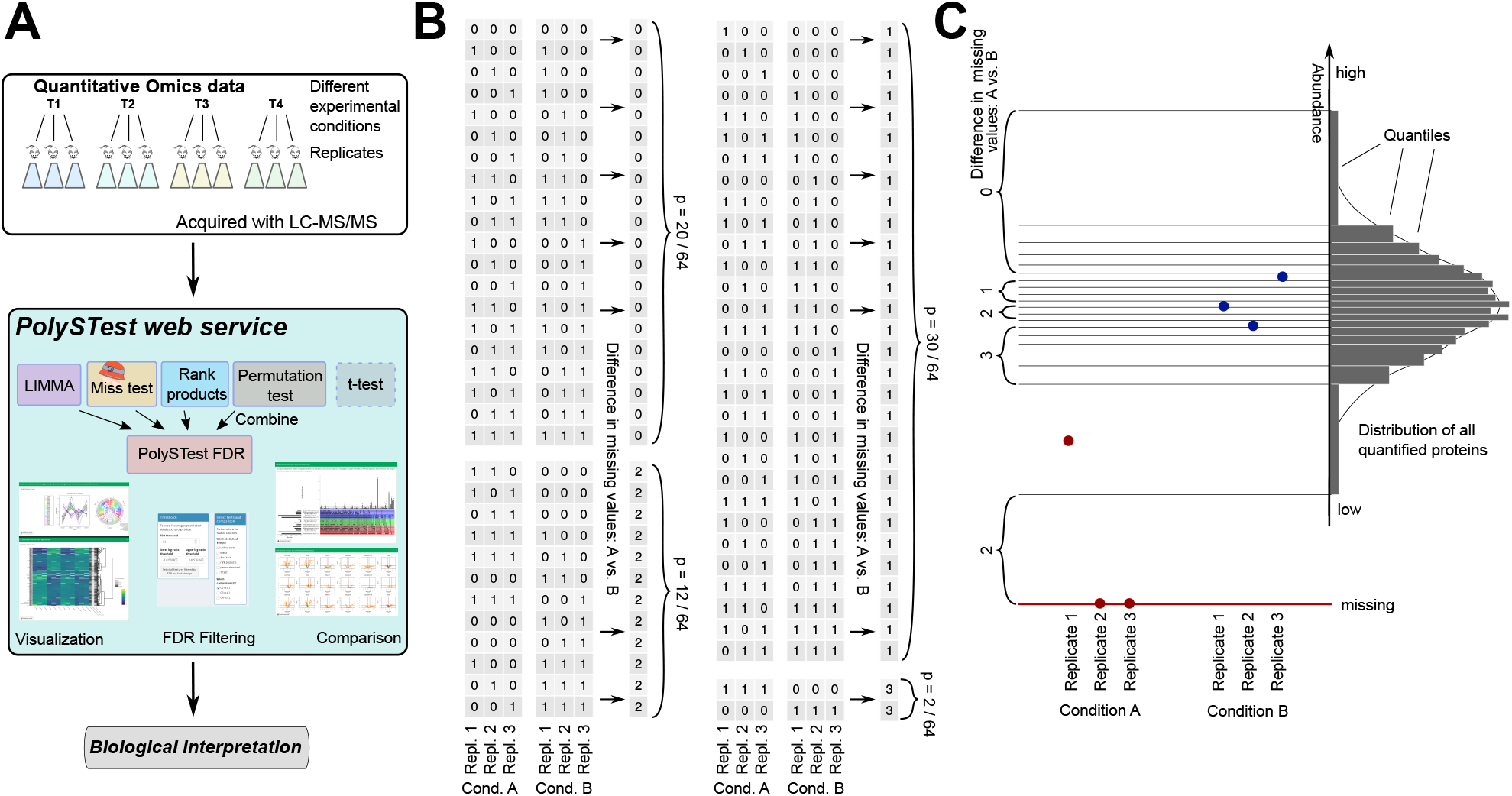
**A** Overview of the PolySTest web service and the used statistical tests. The FDRs from the four statistical tests are combined into a unified PolySTest FDR. **B** Combinations of missing values (zeroes) and present values (ones) in the case of three replicates and two conditions A and B. Given a probability of *p*_NA_ = 1/2 for a missing value, each combination is found with probability 1/64. The numbers on the right count the absolute of the difference in missing values. This leads to p-values of 5/16, 15/32, 3/16 and 1/32 for finding a difference of missing values of 0,1,2 and 3, respectively. **C** Scheme of Miss test on an example of two conditions with three replicates where two of the values of condition A are absent. The distribution of all quantifications in the dataset defines the quantiles used as thresholds for iteratively removing values from the dataset. The differences in the number of missing values are then calculated for each threshold. Note that a different threshold also implies changing *p*_NA_.

### Miss test: A novel statistical test for data with missing values

Miss test integrates missingness with measured protein abundance by reapplying a binary statistics method over a range of different data representations.

To describe the impact of the missing values, we derived the probabilities for each combination of missing values being distributed over the replicates of the experimental conditions. We illustrate the calculation by a simple scenario with 50% of missing values and 3 replicates per each of 2 experimental conditions A and B. The probability for observing a missing value is assumed to be *p*_NA_ = 0.5, which leads to a probability of 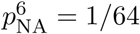 for each combination of missing/non-missing values across the four values (Fig. 1B). We then count the number of cases where the number of missing values in both conditions differs by 0, 1, 2 and 3. A difference of 3 missing values corresponds to only missing values in condition 1 or only in condition 2, thus giving a probability of 2 ⋅ 1/64 = 1/32. A difference of 0, 1 or 2 missing values between the conditions occurs in 20, 30 and 12 different cases. Hence, the probability to find a difference of 0, 1, 2 and 3 missing values is given by the probabilities 5/16, 15/32, 3/16 and 1/32, respectively. Finding only missing values in one condition would then correspond to a p-value of 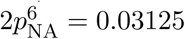.

This calculation only accounts for the presence of missing values. Quantitative values are taken into account by repeating the above-described procedure while artificially increasing the fraction of missing values, *p*_NA_. Our method iteratively reduces the quantitative data by removing low abundant values for a set of 100 abundance thresholds which are calculated from the distribution of the full dataset. The newly introduced missing values of the reduced dataset are therefore abundant dependent. This approach enforces more significance to proteins with high abundance in the condition where less values are missing and where the other condition consists of missing and low abundant protein values. Figure 1C shows an example where all values of condition B are higher than the one value of condition A. This leads to a maximum difference of three missing values. If the one value of condition A had higher abundance than the lowest value of condition B, then the iterative process would only find a maximum difference of 2 missing values, and thus report a higher p-value, i.e. lower significance. The overall performance of Miss test will be thoroughly tested in the next section.

### Performance of Miss test

Here we assess whether Miss test follows the distributions of a common statistical test and whether it can rescue features with many missing values that otherwise would be discarded.

#### ”Empty” data

Purely random data corresponds to a global null hypothesis and should result in a uniform p-value reference distribution. This is crucial to directly compare Miss test to other tests and to ensure that common methods for correction for multiple testing can be used. As the p-values of interest can be very small, we plotted the empirical cumulative distribution of the p-values for normally distributed artificial data sets simulating a variety of scenarios with missing values. Different types of missingness were achieved by setting different weights with regard to protein abundance for the random removal of values in the full artificial data set. This *abundance-dependence* was varied between purely random removal (*µ* = 0) and strong preference for low abundant values (*µ* = 100).

Suppl. Fig. S1 shows that the p-value distribution stays approximately uniform for different replicate numbers. We confirmed this for cases where we varied the fraction of missing values and their abundance-dependence. Suppl. Fig. S2 shows that the p-values still are uniformly distributed when decreasing the number of proteins. We also counted the number of proteins with FDR*<*0.1. For all tests comprising different replicates, 1,000 or 10,000 features, maximally 1 false positive was found, with an overall frequency of less than 5% of all 192 tested data sets which is well below the expected 10%. This confirms absence of false positives in purely random data.

#### Data with ground truth

Next, we investigated how well Miss test recognizes regulated features in artificial data with ground truth. We took data identical to the “empty” random data and added an offset to a fraction of the features to simulate their differential regulation. Thus we divided the data into positives (displaced features) and negatives (random features) which allows validating the results of the statistical tests.

Fig. 2A shows a typical case of noisy data consisting of 1,000 features including 50 features that were different by having only slightly shifted values to either side (*δ* = 1.5) compared to the simulated variance within replicates. Miss tests detects 24 DRFs (FDR*<*0.05). To check how this number is influenced by missing values, 20% of all values were removed assuming a light abundance-dependence on the values via weighted removal (Fig. 2B). Miss test still found 14 of the 50 features to be differentially regulated between both simulated conditions, including cases with few missing values per feature. Reduction to half of the data decreased the number of DRFs to 8 (50% missing values, Fig. 2C). The majority of these features would not have been tested by a common statistical test as only 0-1 values were available in one condition. The performance of Miss test improved with more statistical power (10 replicates per condition, Fig. 2D) or having a stronger relation between missing values and their original abundance (abundance-dependence *µ* = 10, Fig. 2E). In all cases, the number of false positives was kept within the desired range of 5%. Miss test correctly identifies DRFs and becomes powerful for data with many missing values by rescuing otherwise untested differentially regulated proteins.

**Figure 2:**
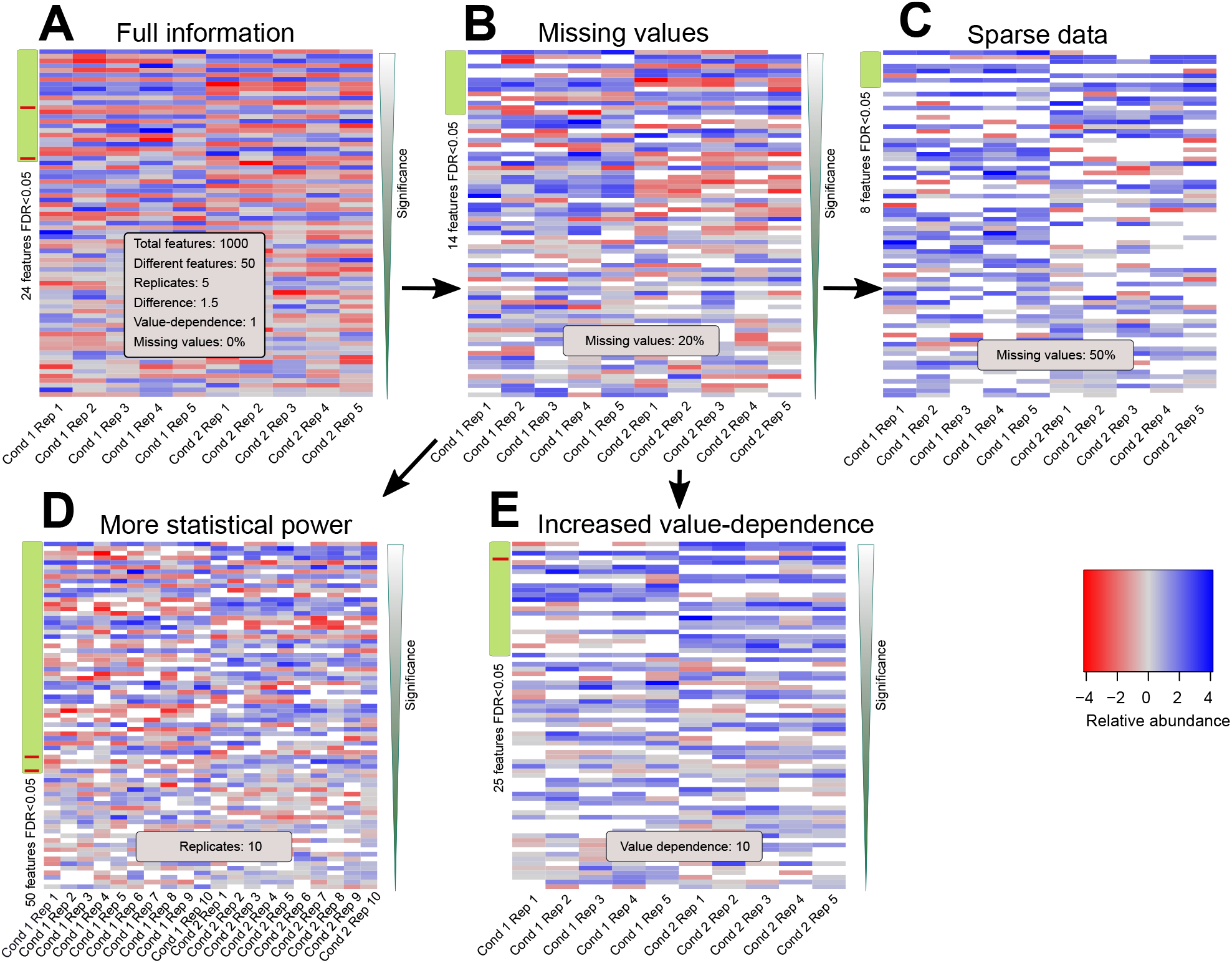
Performance of Miss test for different artificial data sets with missing values. **A-E** Most significantly changing features according to Miss test for differently parametrized artificial data sets. True and false positives are indicated by green and red sidebar values, respectively.

### Performance and complementarity in comparison with other statistical tests

In order to find out whether Miss test is sufficiently powerful as stand-alone test and how it complements standard statistical tests by rescuing otherwise lost proteins, we compared the performance of the Miss test with the performance of the four commonly used statistical tests LIMMA, rank products, permutation test and t-test. To describe and employ their complementary power, a *PolySTest FDR* was calculated combining the FDRs of all used statistical tests except of the t-test: each FDR from LIMMA, Miss test, rank products and permutation test was corrected for multiple testing by the method of Hommel [22] and then the smallest value was taken.

For an overall comparison, we used ROC curves to assess the performance of the different tests applied to hundreds of artificial data sets (see Supplementary Fig. S3 for an example). ROC curves and AUCs (area under curve), despite of being a widely used method to compare test performance, were not able to accurately describe test performance in many cases due to the rather jumpy nature of true and false positive rates, to sometimes low numbers of DRFs, and more importantly to the regions of interest being in the very low range of false positive rates.

Hence, test performance was additionally investigated by means of the following validation criteria at a fixed FDR threshold: (i) sensitivity (or yield) describing the number of detected DRFs, i.e. features that were found to be differentially regulated, (ii) confidence of the found DRFs given by the *true FDR* which was calculated from the number for true positives (TP) and false positives (FP), *tF DR* = *F P/*(*F P* +*T P*), and (iii) comparison of the given FDR threshold to the true FDR to validate how well the test predicts the fraction of false positives. Given these criteria, an optimally performing test would yield a high number of DRFs, keep the true FDR below the FDR threshold but does not underestimate the false discovery rate too drastically.

At an FDR threshold of 0.01, LIMMA was most sensitive at low replicate numbers and low or absent abundance-dependence of the missing values while controlling the false positives at an acceptable level (Fig. 3A). Remarkably, PolySTest maintained a similar number of differentially regulated features but reduced the amount of false positives. Miss test performance improved when increasing the number of replicates or having many missing values. For five replicates (Fig. 3B), most DRFs were detected by LIMMA but PolySTest showed greater sensitivity and confidence through the rescue of additional proteins by Miss test. Higher abundance-dependence of missingness and sparser data sets prevented common statistical tests from calculating p-values because no variance estimate can be calculated from 0-1 values alone. In this case (Fig. 3C), Miss test greatly outperformed the other tests. In all cases, PolySTest kept the true FDR below 0.01, thus correcting cases where one or more of the tests overestimated the fraction of false positives.

**Figure 3:**
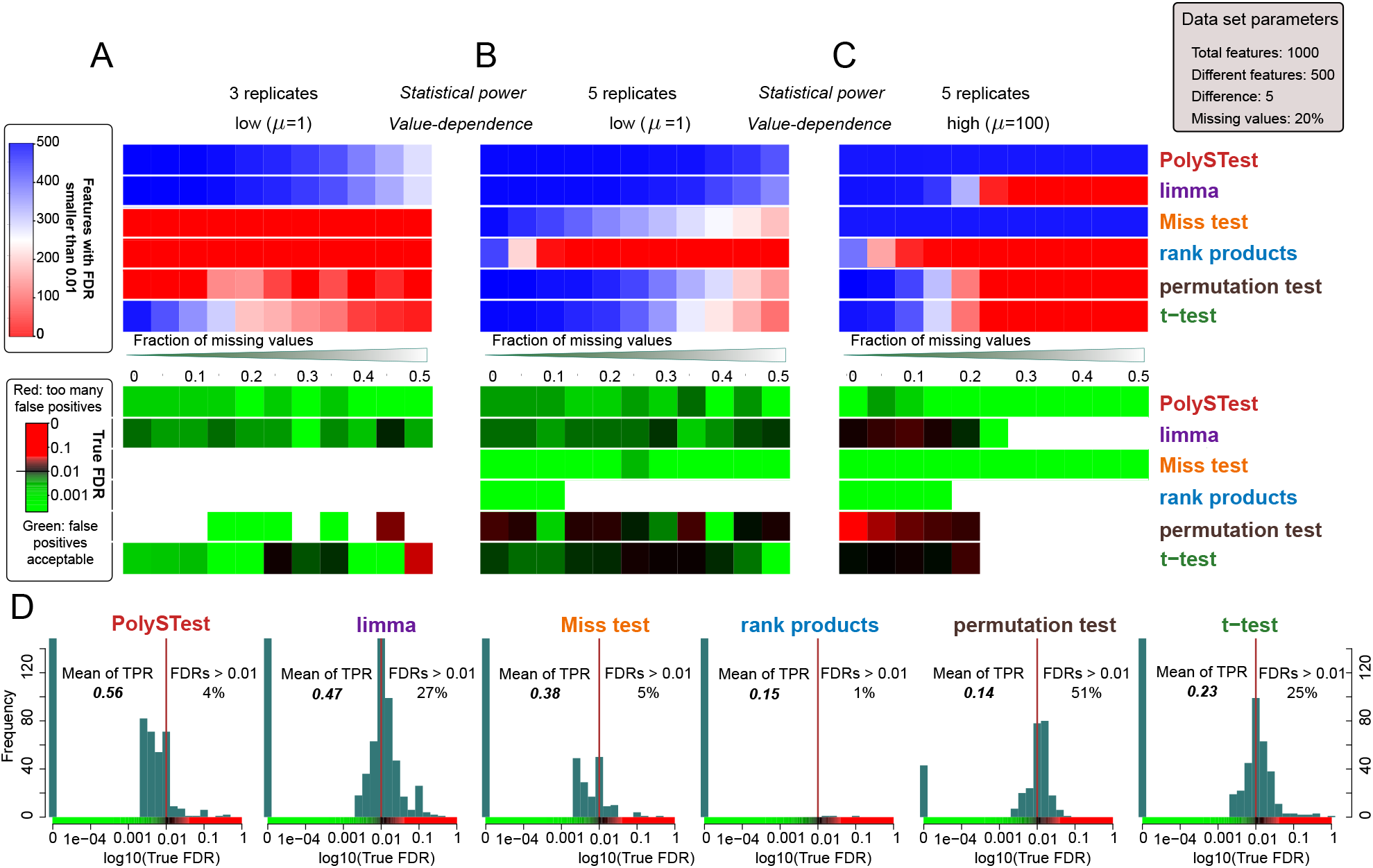
Complementarity of the different statistical tests improves DRF detection. **A-C** Number of DRFs with given FDR smaller than 0.01 and calculated true FDR (from ground truth) for three representative artificial data sets. The results were assessed for different amounts of missing values and show that Miss test complements and therefore improves both DRF detection and confidence for sufficiently high replicate numbers and abundance-dependence of missing values. **D** Distribution of true FDRs at a given FDR of 0.01 for 1,584 artificial data sets spanning 3-10 replicates, 2-100 truly regulated proteins shifted by 1, 1.5, 2 and 5 from their original random value, and 0-100 abundance-dependence *µ* and missing values of 0%-50%. This shows that combining the different statistical tests provide optimal numbers of true positives with high confidence (low percentage of too high true FDRs) over the range of tested data sets.

By thorough comparison of the statistical tests over all artificial data sets, we assessed overall sensitivity (mean of all true positive rates, here denoted mean of TPR) of each test and the distributions of the measured true FDRs (Fig. 3D). This confirmed LIMMA to produce high yields of differentially regulated features at acceptable amounts of false positives. The rank product test showed very high confidence results but suffered from low DRF numbers. Permutation test and t-test showed a low number of DRFs with often too high numbers of false positives. In comparison, Miss test depicted the second highest mean true positive rate and high confidence. As main result, the PolySTest was most accurate in estimating the false discovery rate and produced the largest yield of differentially regulated features. Hence, the considered different statistical tests increase their power when used in a complementary way. We could confirm these general insights for different total numbers of features 500 and 5,000 (Suppl. Figs. S4 and S5).

### PolySTest boosts detection of differentially regulated features in experimental data

We evaluated the performance of the different statistical tests on two published experimental data sets, here denoted *3-species* and *2-species*, with ground truth created by mixing the extracts of different species with pre-determined ratios.

The *3-species* data set [12] contains protein measurements of *E. coli*, yeast and human samples where the former two were mixed at different concentrations keeping the concentration of human proteins constant. By using a mixture of three, one can avoid strong bias towards one side, which can introduce large numbers of false positives as common normalization techniques start to fail. Protein concentrations were measured using SWATH technology [23].

The *2-species* data set [13] contains an *E. coli* extract mixed with a HeLa extract at four different ratios. Protein abundance was measured using label-free quantification on a quadrupole orbitrap mass spectrometer which is considered to have different behavior compared to SWATH given the distinct data acquisition. This different MS platform should also result in a higher abundance-dependence of missing data.

Supplementary Table S1 summarizes the number of true and false positives at a FDR threshold of 0.01 for both ground truth data sets. LIMMA and PolySTest produced the highest sensitivity. Similar to our observation for the artificial data, PolySTest reduced the number of false positives by up to 50%.

When comparing the different tests applied to the *3-species* data on a ROC curve (Fig. 4A), rank products seems to outperform both LIMMA and PolySTest. However, rank products becomes far more conservative than the other tests and produces only very few DRFs for commonly used FDR thresholds (for example FDR 0.01 in Fig. 4B). We additionally inspected sensitivity and confidence and compared the true FDR to the given threshold (Fig. 4C). PolySTest provided accurate confidence thresholds at the price of a slightly lower sensitivity. Nevertheless, most tests overestimate the false positives and we will address to explain this observation in the following section.

**Figure 4:**
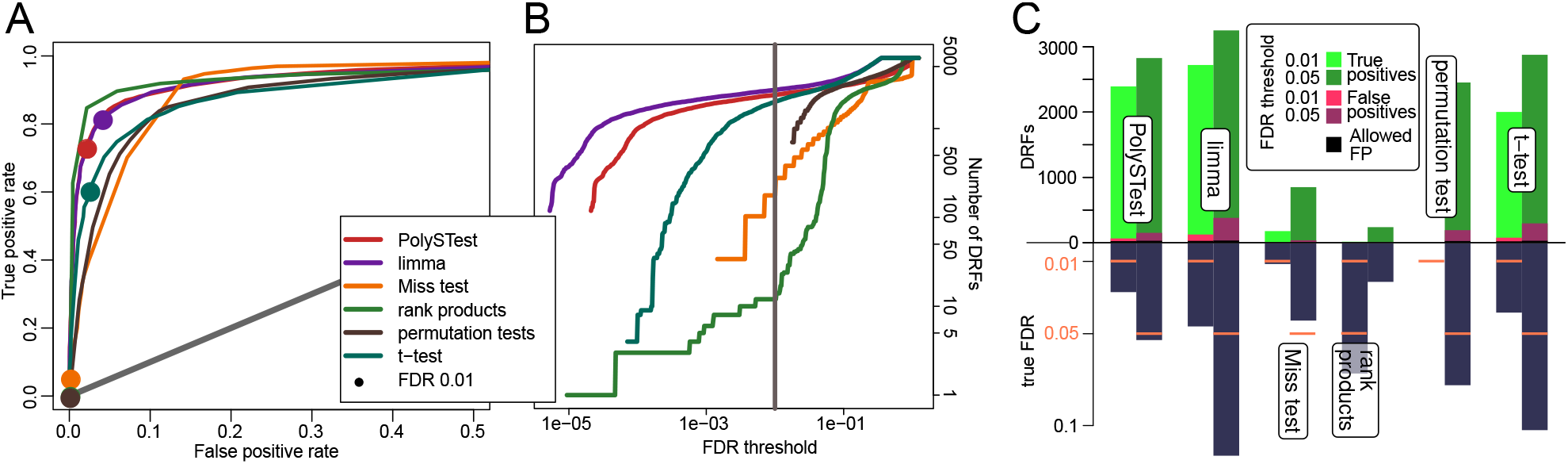
PolySTest provides a high number of true positives at typical confidence levels for the *3-species* data set. **A** ROC curve comparing the different statistical tests with filled circles indicating an FDR of 0.01. Rank products seems to perform best but produces only a very low number of DRFs (**B**). **C** Comparison of test performance on basis of sensitivity and confidence for FDR thresholds of 1% and 5%.

The *2-species* data extends the comparison to four conditions with increasing concentrations of the added *E. coli* extract. Here, the analysis suffers from differential bias as all *E. coli* proteins have higher concentrations when compared to the first condition used as reference. This bias has strong impact on the normalization and we therefore did re-normalize the quantitative data using the cyclic loess method from the LIMMA package.

Given that the data was acquired with the label-free approach, missing values were likely to come from low abundant proteins for which the peptide peaks did not pass the detection limit. This higher abundance-dependence leads to a higher sensitivity of PolySTest visible by a considerable increase of the ROC curve of the PolySTest score when comparing it to the other statistical tests (Fig. 5A). This increase in performance results benefits from proteins that were completely absent in the first condition and therefore were only considered by the Miss test.

**Figure 5:**
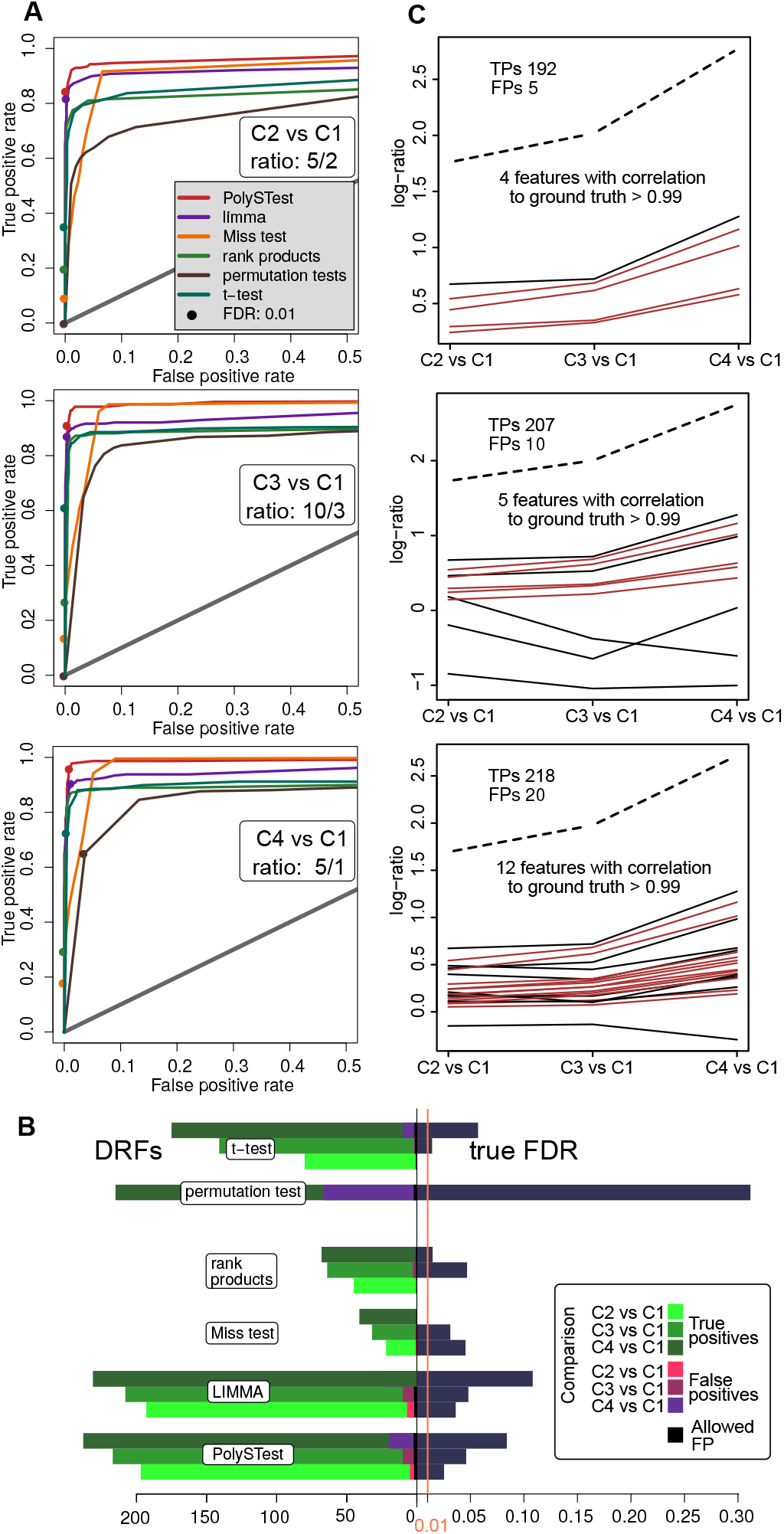
PolySTest outperforms individual tests for the 2-species data set as Miss test adds significant proteins. **A** ROC curves show that combined tests, mostly supplied by the Miss test, considerably increase the true positive rate. Filled circles correspond to an FDR of 0.01. **B** Comparison of test performance on basis of sensitivity and confidence (FDR: 0.01). **C** Higher concentrations of E. coli proteins in mixture leads to more false positives which however follow the trend of the mixture as confirmed by high correlations to the given ratios. The dashed line corresponds to the averaged log-ratios of all *E. coli* proteins. Red lines are proteins with correlation *>* 0.99.10/3

Consistently with the previous results, PolySTest decreased the number of false positives when compared with LIMMA (Fig. 5B). However, the test still overestimated the FDR and this effect increased with the concentration of the *E. coli* extract.

In summary, combining the different statistical tests to a unified PolySTest score resulted in higher confidence at either low cost of sensitivity or even its increase by rescuing significant proteins with many missing values.

### High number of false positives explained by wrong peptide identifications in MS data

In contrast to the results for the artificial data, here most statistical tests suffered from too many false positives. We investigated whether the false positives could result from artifacts in the upstream data analysis pipeline, for example due to wrong peptides identifications that assign a peptide from the wrong species.

To get a rough estimation of wrong peptide identifications within the false positives in the *3-species data* set, we counted how many peptides per protein determine their identity. Figure S6 shows that the majority of false positive proteins is based on 1-2 peptides which is different from the peptide counts in the full dataset where higher peptide counts are much more frequent. Hence, we can assume a higher number of false identifications within them and therefore these human proteins might be derived from actual differentially regulated proteins from either *E. coli* or yeast. More accurate true FDRs might be achieved by removing proteins that were identified by only one peptide.

In the *2-species* data, we took advantage of the multi-dimensional setup comparing 4 different ratios between *E. Coli* and human samples. This allowed to look for proteins with potentially wrong identifications by comparing their expression profile over all five *E. coli* concentrations (Fig. 5C). Then, the protein expression profiles of wrongly assigned false positives will correlate highly with the overall profile of the *E. coli* extracts shown as dashed line. For that, we calculated Pearson’s correlation between the false positives and the mean of the ratios of the true positives. The resulting ratios were lower but still many proteins correlated strongly (*>*0.99) with the true positives. This most likely comes from human proteins, for which a fraction of *E. coli* peptides was wrongly identified as human peptides.

Our analysis shows that too tolerant thresholds for the peptide identification can have major impact on the down-stream statistical analysis by assigning the wrong protein to a DRF. On the other hand, we suspect that the number of wrong identifications leading to false positives is lower in real biological samples where biological variation should suppress these false hits more efficiently.

### Higher coverage of biologically relevant proteins

After comparing the performance of the different tests on ground truth data, we applied PolySTest on biological data from experiments where cells undergo drastic changes. Immortalized human satellite cells (KM155C25) [15] were differentiated into muscle cells and samples were taken over 6 time points in biological triplicates: proliferating activated myoblasts (Day −1) and 5 days following the initiation of differentiation into mature myocytes (Day 0, 1, 2, 3 and 4). The data was acquired using an Orbitrap Fusion™Lumos™Tribrid™Mass Spectrometer and analysed in MaxQuant using chromatography alignment feature and label free quantitation (LFQ) of proteins. This setup allows the evaluation of the result on biological grounds, i.e. by exposing the principal biological pathways known to change during differentiation, particularly pathways that are specific for muscle cells and cell cycle.

At an FDR threshold of 0.05, LIMMA detected the highest number of DRFs (2,620 proteins, day 4 versus day −1) followed by PolySTest (2,294) (see Suppl. Fig. S7 for details). We investigated how well this larger number of DRFs described the biological content by assessing the relevant biological pathways expected to be altered during muscle differentiation. We first focused on the Striated Muscle Contraction Pathway (wikipathways ID WP383) where we quantified 27 out of 38 proteins (Fig. 6A). When comparing the latest differentiation state to the satellite cells, PolySTest detected a higher number of 26 proteins in this muscle-specific pathway compared with 19 significant proteins detected by LIMMA (Fig. 6B, “C6 vs C1”). This higher number resulted mostly from Miss test which supplemented proteins that were absent or nearly absent in the first condition (Fig. 6C). PolySTest increased the number of DRF proteins in the following pathways, known to be altered during muscle differentiation: Arrhythmogenic Right Ventricular Cardiomyopathy (24 versus 20 proteins), G1 to S cell cycle control (19 versus 15 proteins) and Cell Cycle (29 versus 26).

**Figure 6:**
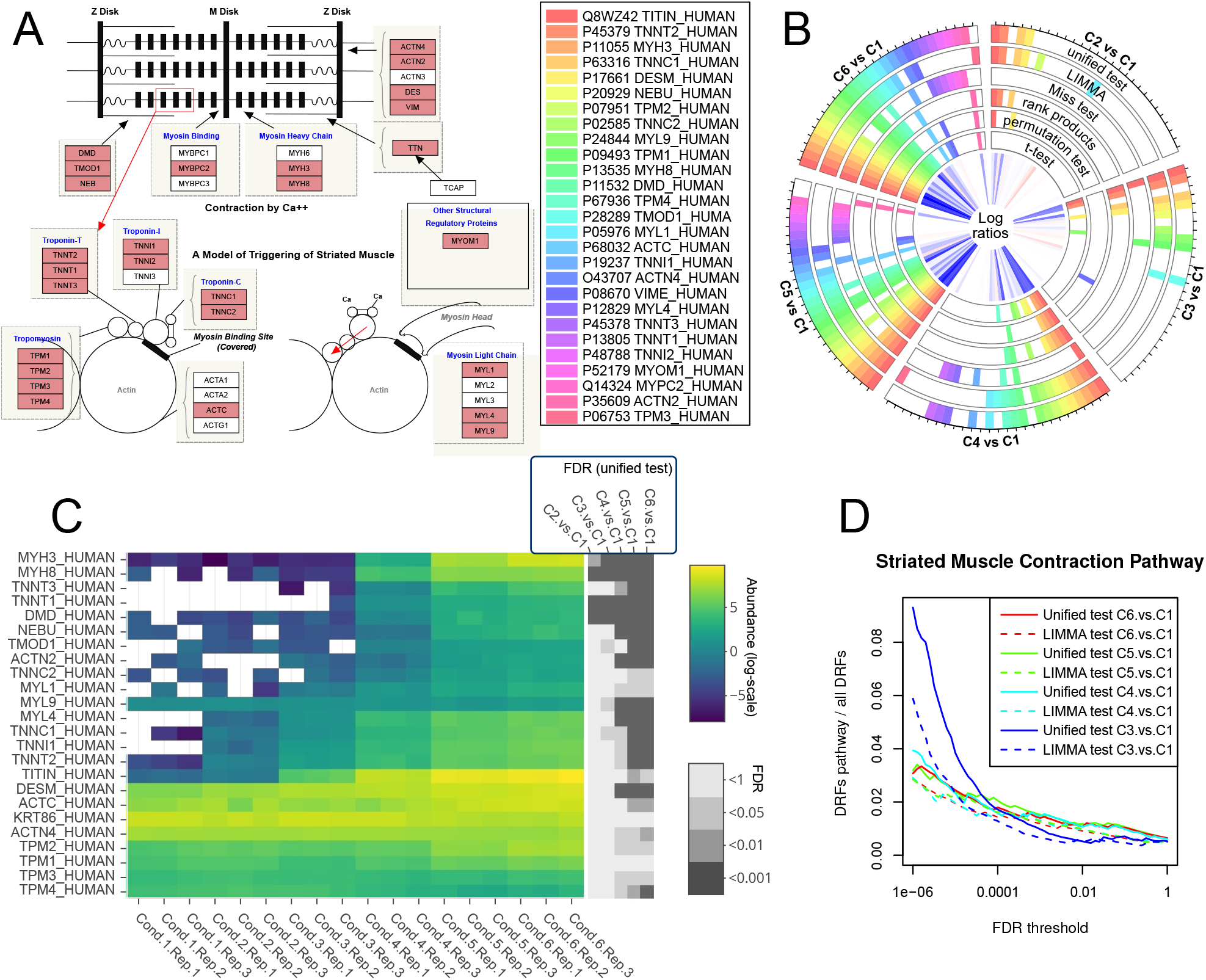
Improved recognition of proteins from Striated Muscle Contraction Pathway (Wikipathways) when differentiating into muscle cells. **A** Illustration of the pathway and all 27 quantified proteins (red). **B** Comparison of the significantly changing proteins of the different statistical tests for an FDR threshold of 0.05. A filled box denotes a protein with FDR *<* 0.05. **C** Unscaled abundance changes and calculated false discovery rates for the different comparisons (3 proteins with more than 50% missing values are not shown in this figure). **D** Enrichment of differentially regulated pathway proteins versus all DRFs comparing PolySTest and LIMMA.

Proteins detected by PolySTest alone but not LIMMA were submitted to pathway enrichment analysis and could be assigned to muscle-related Reactome and Wikipathways pathways in most cases (Suppl. Fig. S9). For instance, the retinoblastoma gene is known to play a crucial role in muscle differentiation [24]. This confirms that Miss test and PolySTest contribute with relevant biological information. Proteins identified by LIMMA and but not PolySTest showed enrichment of more general pathways such as translational control and mRNA processing.

We tested robustness of the enriched Striated muscle contraction pathway with respect to changes of the FDR threshold (Fig. 6D). When measuring the fraction of DRFs in the pathway compared to their total number, we observed a decrease when increasing the FDR threshold, which is consistent with additional, less specific pathways entering the picture. PolySTest outperformed LIMMA over the entire FDR range as well as for the different differentiation states.

We now extended the analysis to the most significant pathways and pathways associated with muscle cells and cell cycle. Visual separation of the figures into pathways with a similar decrease for the enrichment for an increasing FDR threshold showed that this behavior was explicitly found for pathways strongly associated with muscle differentiation and cell cycle. Interestingly, PolySTest outperformed LIMMA in all these cases but not the ones of supposingly less or not affected pathways.

## Discussion

Detection of a maximal number of differentially regulated features with correctly assigned false discovery rate is a challenge. We addressed the demand for a better integration of missing values, robustness of the statistical tests and a friendly and interactive user interface (Suppl. Figs. S10 and S11). We have developed a software tool for in-depth statistical testing of quantitative omics data which we evaluate extensively on data from proteomics experiments. The interactive web interface runs the five statistical tests moderated t-test (LIMMA package), rank products, permutation test, t-test and Miss test where the latter was newly implemented. Complementarity of the different tests was taken into account by the PolySTest FDR which unifies the FDRs from all tests but the t-test to a high-confidence score as will be shown below. The test results can be visually compared and further assessed via an interactive table and down-stream visualization in terms of p-value histograms, volcano plots, heatmaps, expression profiles, upset plots and a circular graphics to compare the different statistical tests.

PolySTest detected differentially regulated features with higher confidence and similar or higher sensitivity when compared to single tests such as moderated t-test (LIMMA). The new Miss test was designed to not require a strong hypothesis for the missingness of the values. In comparison to prior approaches including missing features, it also values feature abundance levels. We found that Miss test provides important complementary information to increase sensitivity of statistical tests in data with missing values. In addition, this test allows disregarding commonly used value imputation which is very likely to seriously perturb the statistical testing by adding wrong information.

The complementary performance of Miss test, LIMMA, rank products and a permutation test were thoroughly validated on artificial data, experimental data with ground truth and an experimental data set with known regulated biological pathways. The combination of multiple tests was found to be more robust to different data structures and data acquisition methods like DIA and DDA and resulted in a more strict control of the false discovery rate. Hence, this approach achieved higher confidence while at least keeping similar sensitivity.

However, we also found that the differential analysis of MS data suffers from wrong identifications. The commonly used 1% FDR for peptide and protein identification influences the statistical tests by assigning wrong peptide sequences to actually regulated proteins. This can be avoided by lowering the threshold for the identification when aiming for high-confidence results e.g. in clinical applications. This counts especially for the application of statistical tests to data consisting of (modified) peptides. With PolySTest giving better FDR estimates and restraining false positives within a reasonable level, we also recommend its usage to this data type.

Our approach and its user-friendly and accessible implementation facilitates robust statistical testing of proteomics and other quantitative omics data while improving the sensitivity and accuracy.

## Acknowledgments

VS was funded by The Danish Council for Independent Research, the Danish National Research Foundation (DNRF82) and the EU ELIXIR consortium (Danish ELIXIR node). ONJ acknowledges generous financial support from the Danish Council for Research, Natural Sciences (FNU) and VILLUM Center for Bioanalytical Science (VILLUM Foundation). We also thank R. Sprenger for helpful discussions.

## Data availability

All methods are freely available through our web service at http://computproteomics.bmb.sdu.dk/Apps/PolySTest or can be downloaded to be run on a local computer. The full source code is available through https://bitbucket.org/veitveit/polystest.

