## Supplementary tables and figures for "PolySTest: Robust statistical testing of proteomics data with missing values improves detection of biologically relevant features"

| | *Unified test* | *LIMMA* | *Miss test* | *Rank products* | *Permutation test* | *t-test* |
|---|---|---|---|---|---|---|
| | 3-species mixture FDR 0.01 | | | | | |
| *TP* | 2329 | **2597** | 175 | 13 | 0 | 1926 |
| *FP (tFDR)* | 64 (0.03) | 124 (0.05) | 2 (0.01) | 1 (0.07) | 0 | 76 (0.04) |
| | 3-species mixture FDR 0.05 | | | | | |
| *TP* | 2677 | **2872** | 814 | 232 | 2262 | 2582 |
| *FP (tFDR)* | 150 (0.05) | 379 (0.12) | 36 (0.04) | 5 (0.02) | 191 (0.08) | 295 (0.1) |
| | 2-species mixture FDR 0.01 (condition 2 vs. 1) | | | | | |
| *TP* | **192** | 186 | 21 | 45 | 0 | 80 |
| *FP (tFDR)* | 5 (0.03) | 7 (0.04) | 1 (0.05) | 0 (0) | 0 (0) | 0 (0) |
| | 2-species mixture FDR 0.01 (condition 3 vs. 1) | | | | | |
| *TP* | **207** | 198 | 31 | 61 | 0 | 139 |
| *FP (tFDR)* | 10 (0.05) | 10 (0.05) | 1 (0.03) | 3 (0.05) | 0 (0) | 2 (0.01) |
| | 2-species mixture FDR 0.01 (condition 4 vs. 1) | | | | | |
| *TP* | **218** | 206 | 41 | 67 | 152 | 165 |
| *FP (tFDR)* | 20 (0.08) | 25 (0.1) | 0 (0) | 1 (0.01) | 64 (0.3) | 10 (0.06) |

Table S1: Comparison of true and false positives within statistical tests for all ground truth data sets. The 2-species data set been normalized using the cylic loess normalization by LIMMA to decrease effects from bias towards one side. TP: true positives, FP: false positives, tFDR: true FDR.

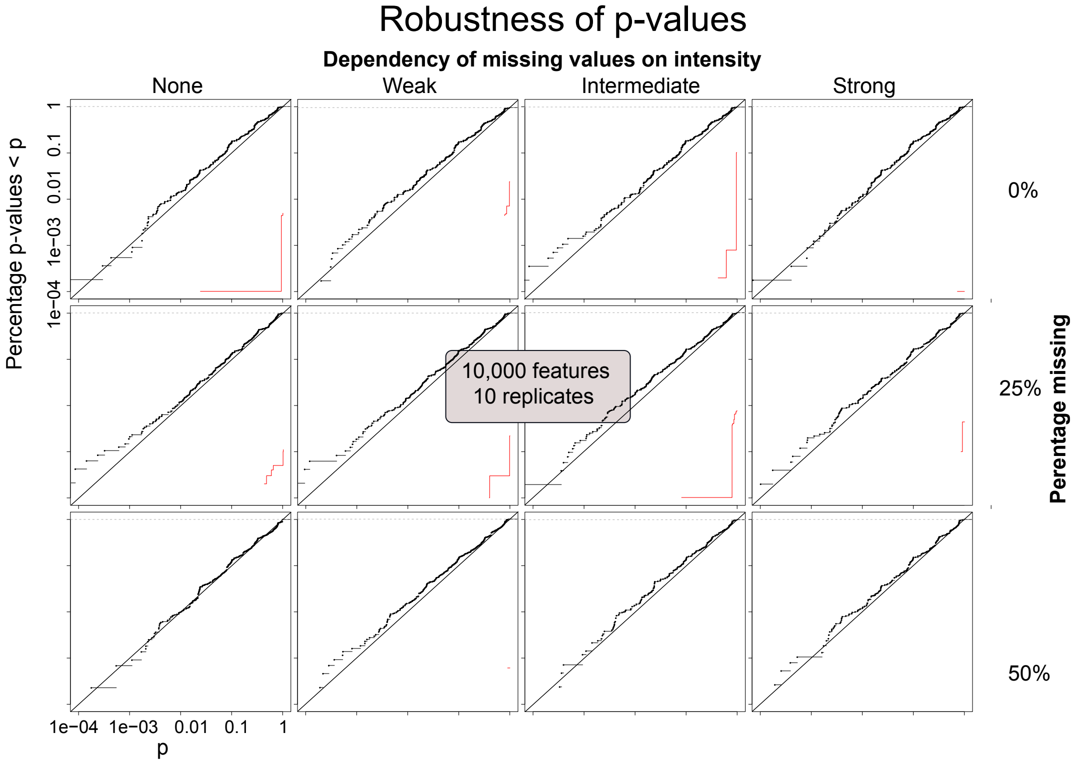


Figure S1: Cumulative distribution of p-values for different amounts and different value-dependent missing values. A linear increase corresponds to an uniform distribution of p-values which is valid down to very low p-values. The red lines denote the fraction of features with an FDR below the give $p$.

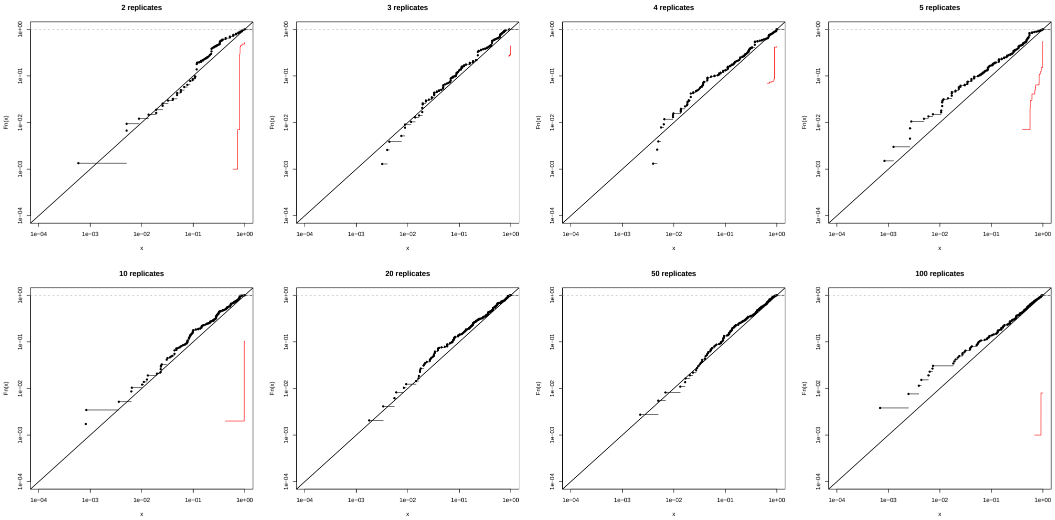


Figure S2: Cumulative distribution of p-values from Miss test for purely random artificial data consisting of 1,000 features.

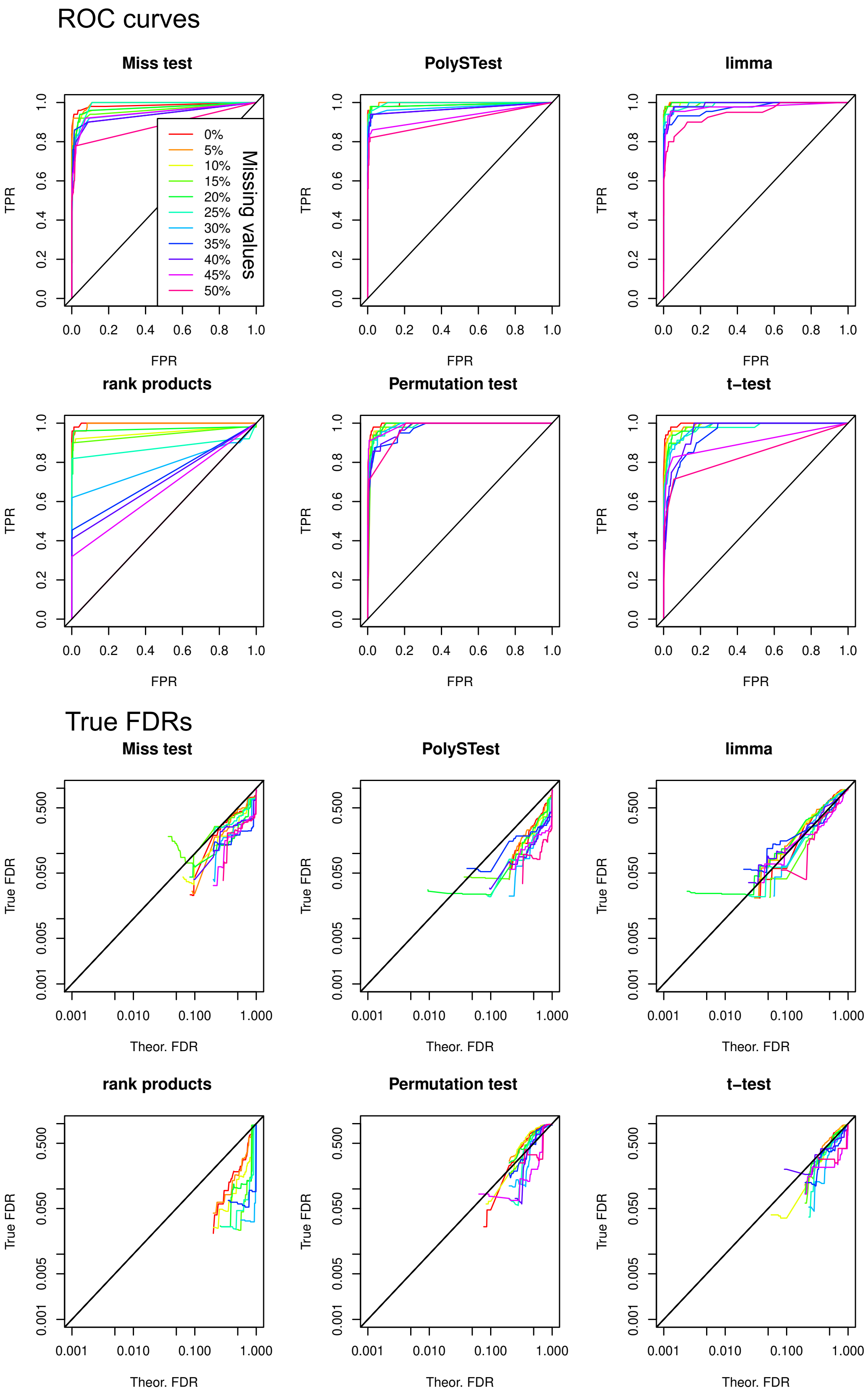


Figure S3: ROC curves and comparison of estimated and real FDR for the data sets from Fig.??C when additionally comparing different percentages of missing values.

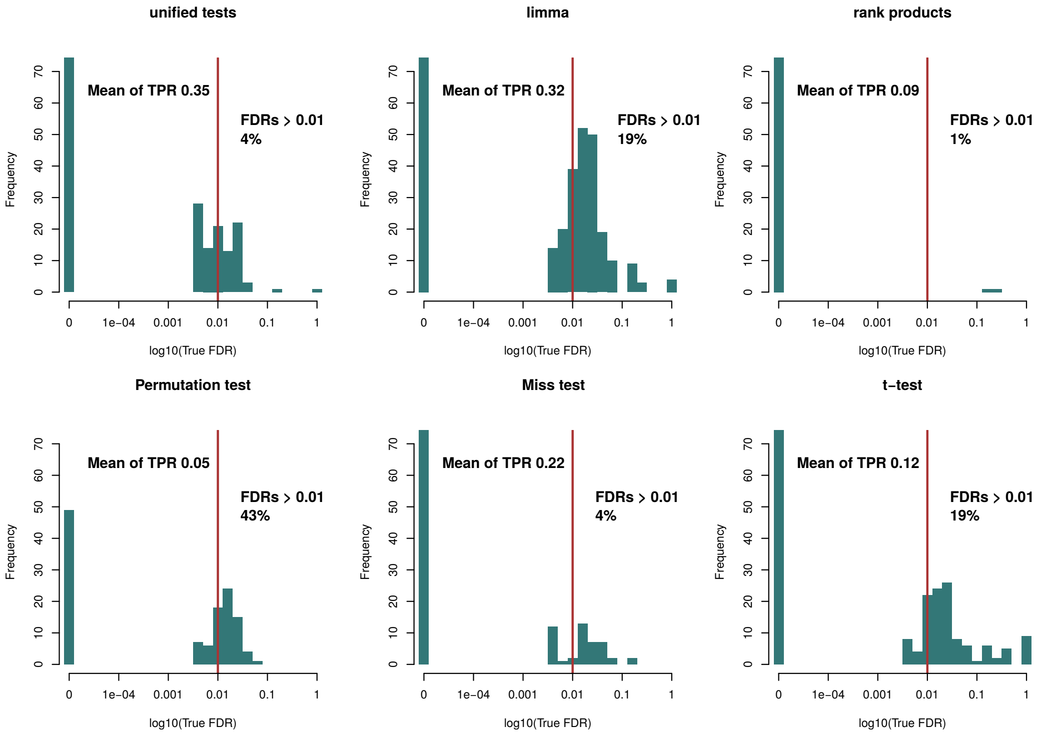


Figure S4: Distribution of true FDRs at a given FDR of 0.01 for 1,584 artificial data sets with 500 features spanning 3-10 replicates, 2-100 truly regulated proteins shifted by 1, 1.5, 2 and 5 from their original random value, and 0-100 value-dependence and missing values of 0%-50%.

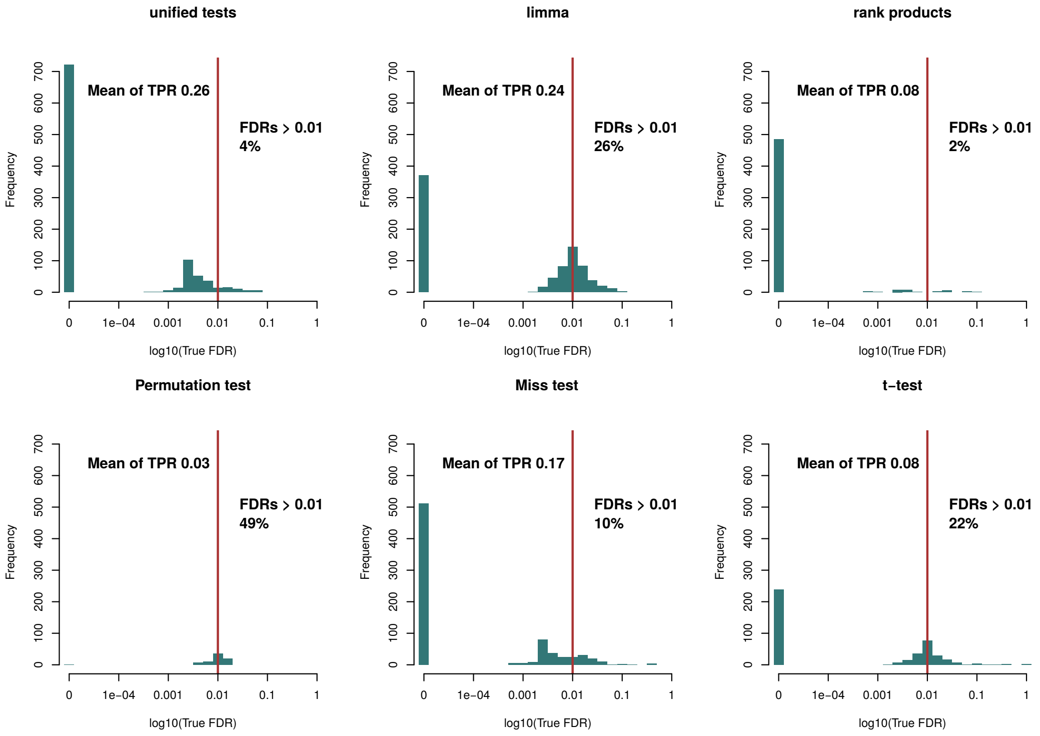


Figure S5: Distribution of true FDRs at a given FDR of 0.01 for 1,584 artificial data sets with 5,000 features spanning 3-10 replicates, 2-100 truly regulated proteins shifted by 1, 1.5, 2 and 5 from their original random value, and 0-100 value-dependence and missing values of 0%-50%.

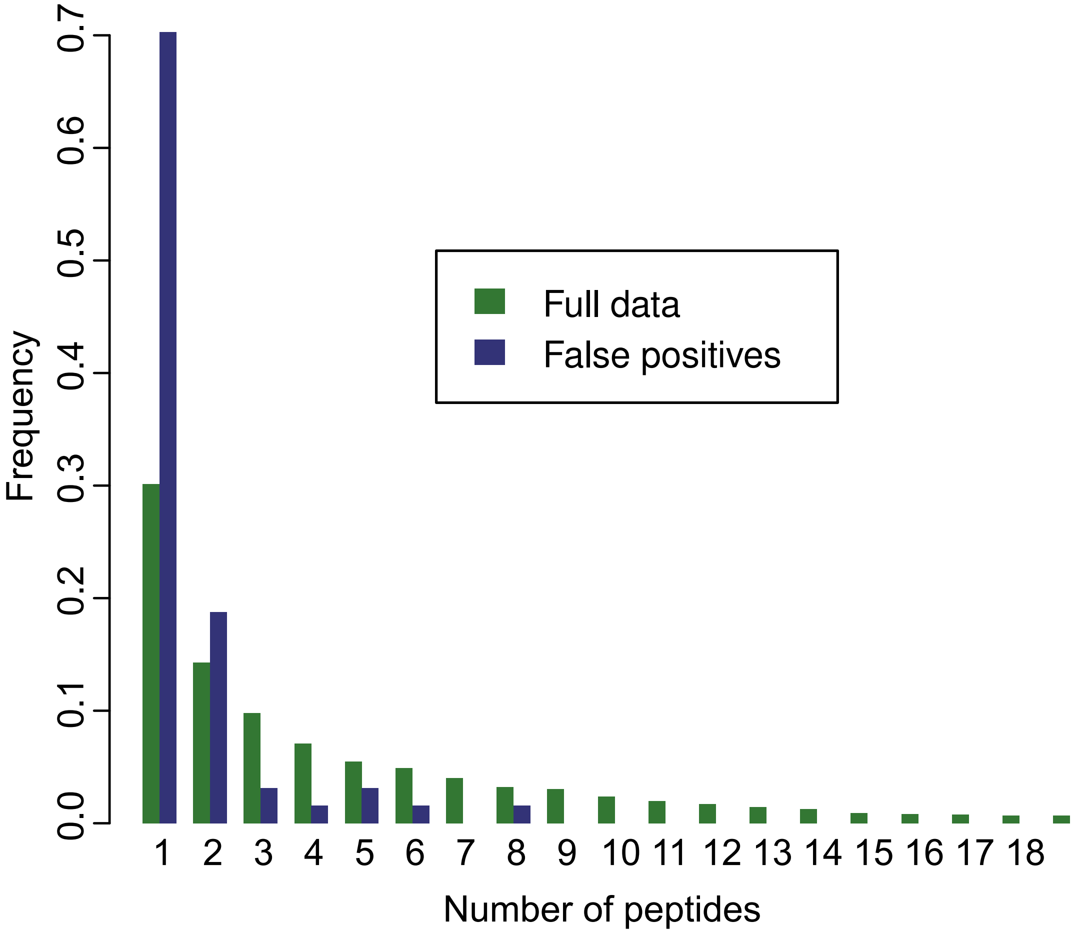


Figure S6: False positives are mostly proteins derived from low peptide numbers compared to the full data set, indicating that they, at least partially, might in fact be wrongly identified true positives.

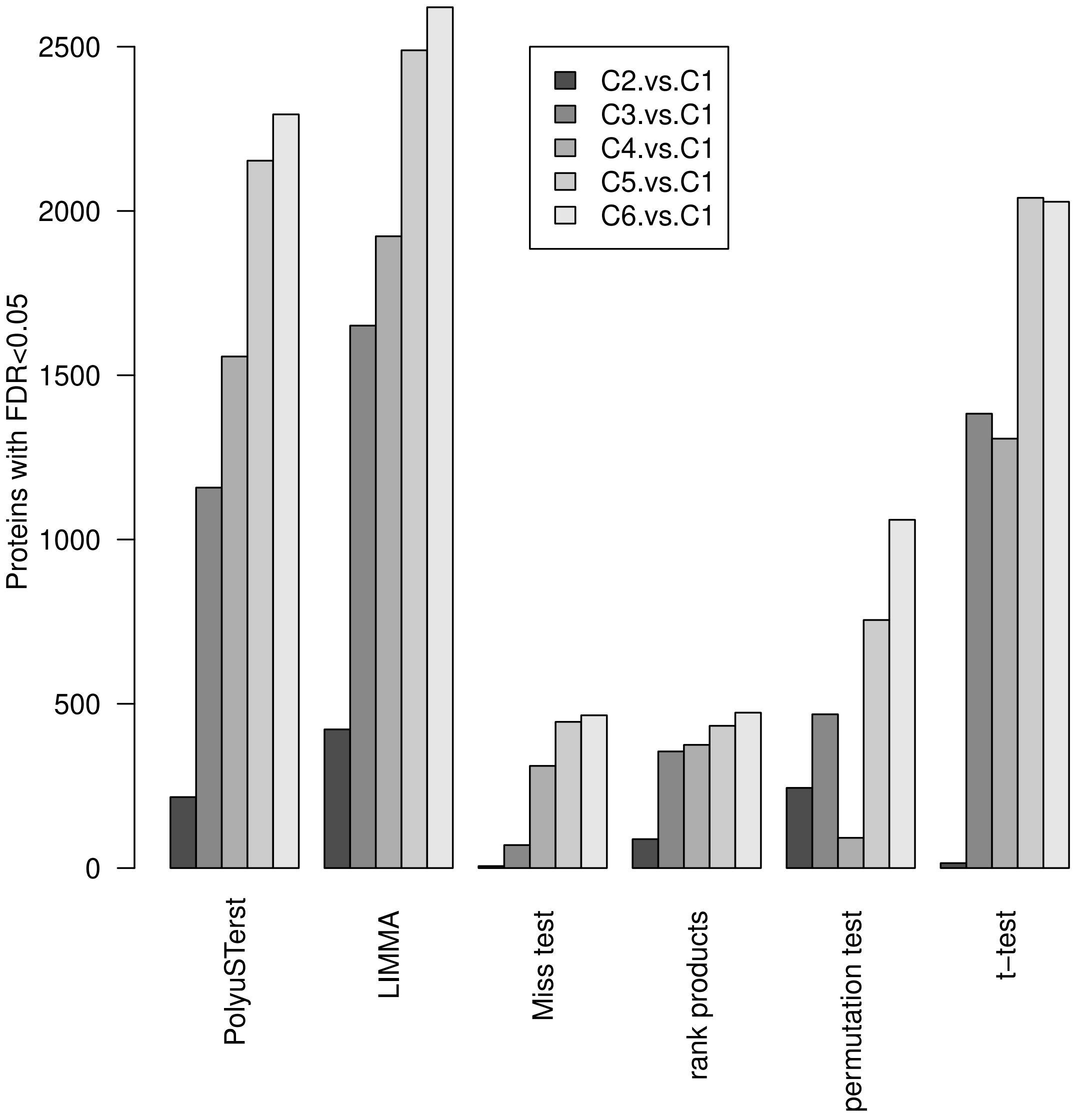


Figure S7: Number of differentially regulated proteins at FDR < 0.05 for the different statistical tests and comparisons of different muscle differentiation states versus stem cells.

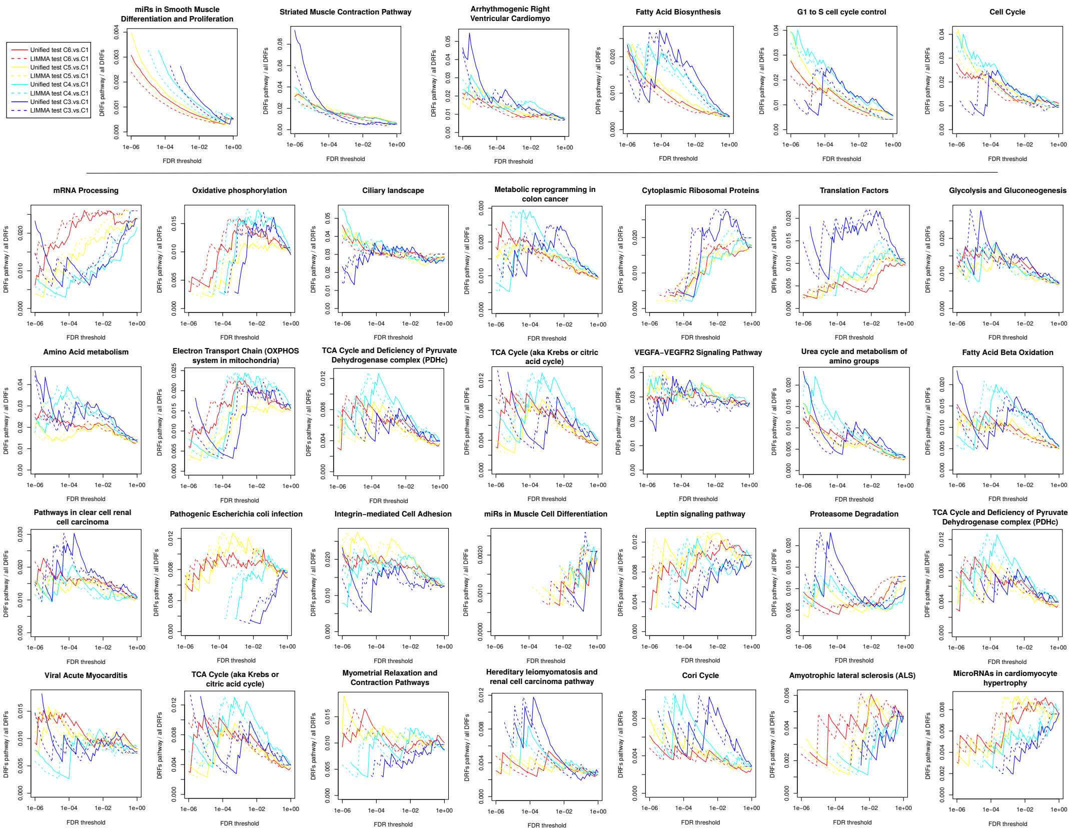


Figure S8: Gene-background ratios of DRFs for most enriched pathways and pathways related to muscle cells and cell cycle. The gene groups were taken from WikiPathways.

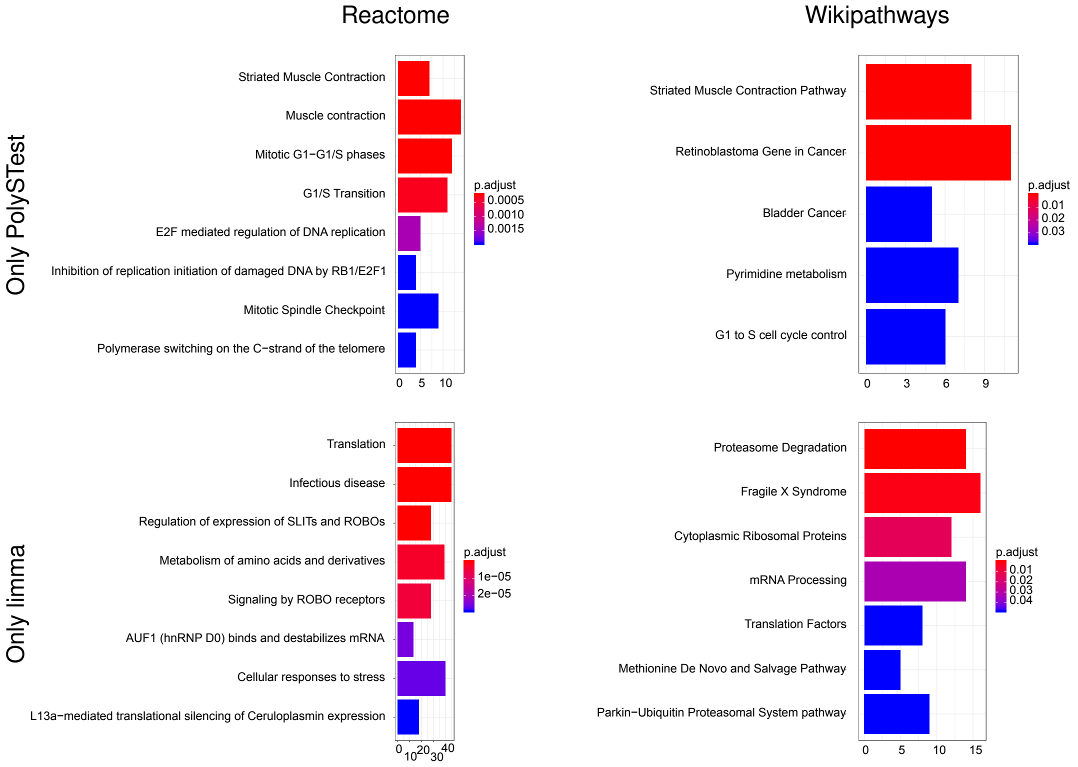


Figure S9: Result from pathway enrichment analysis (Reactome and Wikipathways) for proteins uniquely identified by PolySTest (upper panels) and LIMMA (lower panels). FDR threshold: 0.05.

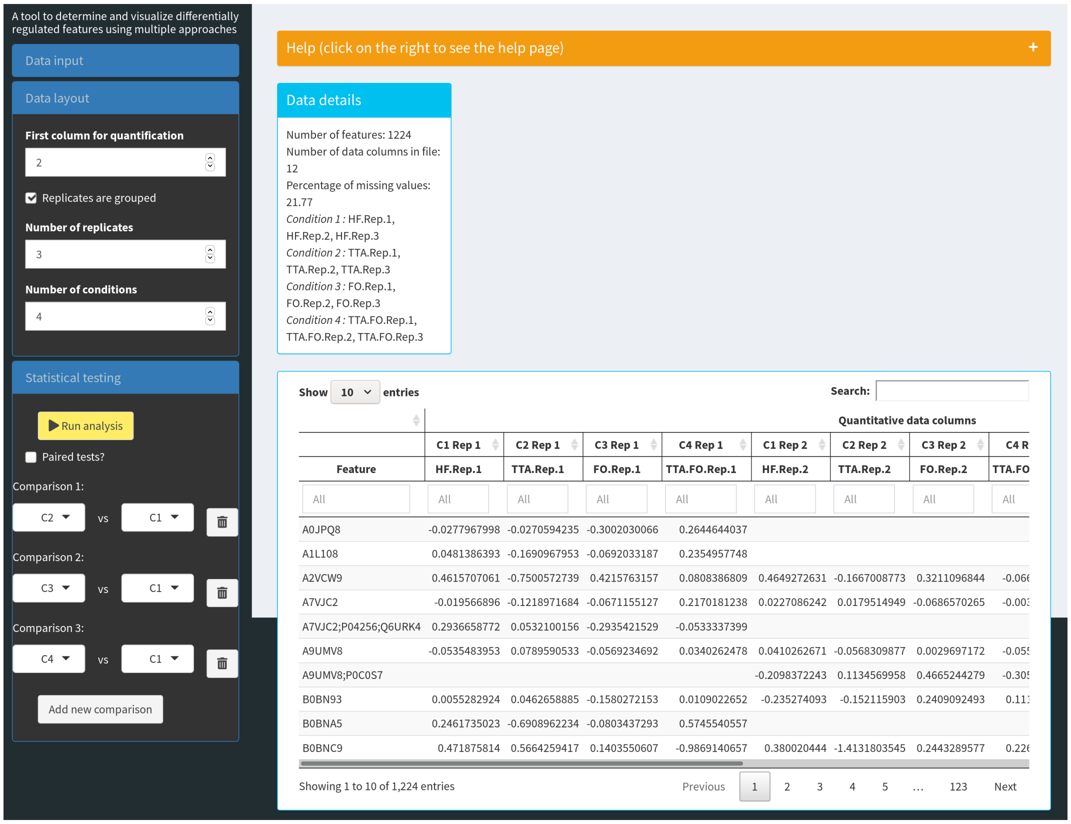

Figure S10: Screenshot of the PolySTest web service. Upload of quantitative proteomics data and setting of experimental design.

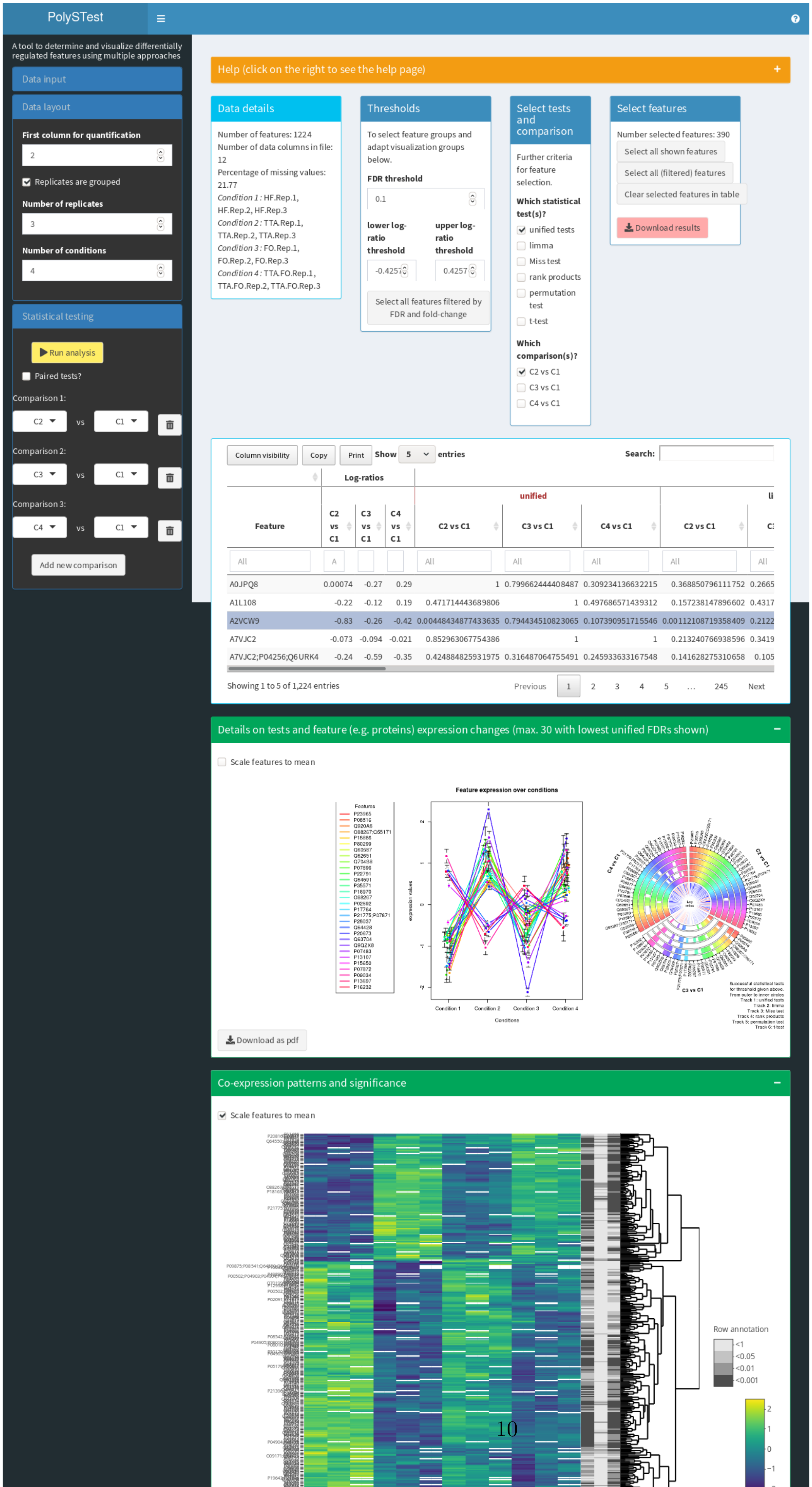
PolySTest
A tool to determine and visualize differentially regulated features using multiple approaches
Data input
Data layout
First column for quantification
2
Replicates are grouped
Number of replicates
3
Number of conditions
4
Statistical testing
Run analysis
Paired tests?
Comparison 1:
C2 vs C1
Comparison 2:
C3 vs C1
Comparison 3:
C4 vs C1
Add new comparison
Help (click on the right to see the help page)
Data details
Number of features: 1224
Number of data columns in file: 12
Percentage of missing values: 21.77
Condition 1 : HF.Rep.1, HF.Rep.2, HF.Rep.3
Condition 2 : TTA.Rep.1, TTA.Rep.2, TTA.Rep.3
Condition 3 : FO.Rep.1, FO.Rep.2, FO.Rep.3
Condition 4 : TTA.FO.Rep.1, TTA.FO.Rep.2, TTA.FO.Rep.3
Thresholds
To select feature groups and adapt visualization groups below.
FDR threshold
0.1
lower log-ratio threshold
-0.4257
upper log-ratio threshold
0.4257
Select all features filtered by FDR and fold-change
Select tests and comparison
Further criteria for feature selection.
Which statistical test(s)?
unified tests
limma
Miss test
rank products
permutation test
t-test
Which comparison(s)?
C2 vs C1
C3 vs C1
C4 vs C1
Select features
Number selected features: 390
Select all shown features
Select all (filtered) features
Clear selected features in table
Download results
Column visibility
Copy
Print
Show 5 entries
Search:
Log-ratios
unified
Feature | C2 vs C1 | C3 vs C1 | C4 vs C1 | C2 vs C1 | C3 vs C1 | C4 vs C1 | C2 vs C1
A0JPQ8 0.00074 -0.27 0.29 1 0.799662444408487 0.309234136632215 0.368850796111752 0.2665
A1L108 -0.22 -0.12 0.19 0.471714443689806 1 0.497686571439312 0.157238147896602 0.4317
A2VCW9 -0.83 -0.26 -0.42 0.0044843487743635 0.794434510823065 0.107390951715546 0.00112108719358409 0.2122
A7VJC2 -0.073 -0.094 -0.021 0.852963067754386 1 1 0.213240766938596 0.3419
A7VJC2;P04256;Q6URK4 -0.24 -0.59 -0.35 0.424884825931975 0.316487064755491 0.245933633167548 0.141628275310658 0.105
Showing 1 to 5 of 1,224 entries
Previous 1 2 3 4 5 … 245 Next
Details on tests and feature (e.g. proteins) expression changes (max. 30 with lowest unified FDRs shown)
Scale features to mean
Feature expression over conditions
Features
expression values
Condition 1
Condition 2
Condition 3
Condition 4
Conditions
C4 vs C1
C2 vs C1
C3 vs C1
Successful statistical tests for threshold given above.
From outer to inner circles
Track 1: unified tests
Track 2: limma
Track 3: Miss test
Track 4: rank products
Track 5: permutation test
Track 6: t-test
Download as pdf
Co-expression patterns and significance
Scale features to mean
Row annotation
<1
<0.05
<0.01
<0.001
